# RBBP6 activates the pre-mRNA 3’-end processing machinery in humans

**DOI:** 10.1101/2021.11.02.466915

**Authors:** Vytaute Boreikaite, Thomas Elliott, Jason Chin, Lori A Passmore

**Affiliations:** MRC Laboratory of Molecular Biology, Cambridge UK

## Abstract

3’-end processing of most human mRNAs is carried out by the cleavage and polyadenylation specificity factor (CPSF; CPF in yeast). Endonucleolytic cleavage of the nascent pre-mRNA defines the 3’-end of the mature transcript, which is important for mRNA localization, translation and stability. Cleavage must therefore be tightly regulated. Here, we reconstitute specific and efficient 3’-endonuclease activity of human CPSF with purified proteins. This requires the seven-subunit CPSF as well as three additional protein factors: cleavage stimulatory factor (CStF), cleavage factor IIm (CFIIm) and, importantly, the multi-domain protein RBBP6. Unlike its yeast homologue Mpe1, which is a stable subunit of CPF, RBBP6 does not copurify with CPSF and is recruited in an RNA-dependent manner. Sequence and mutational analyses suggest that RBBP6 interacts with the WDR33 and CPSF73 subunits of CPSF. Thus, it is likely that the role of RBBP6 is conserved from yeast to human. Overall, our data are consistent with CPSF endonuclease activation and site-specific pre-mRNA cleavage being highly controlled to maintain fidelity in RNA processing.

## Introduction

Eukaryotic protein-coding pre-mRNAs undergo multiple processing steps during transcription by RNA polymerase II. These include 5’-capping, splicing, and 3’-end processing (Hocine et al., 2010). During this latter process, a cleavage event defines the 3’-end of the mature mRNA and is linked to transcription termination (Buratowski, 2005; Liu and Moore, 2021). A poly(A) tail is added to the resultant free 3’-end, marking the mRNA for nuclear export and controlling mRNA stability and translational efficiency in the cytoplasm (Passmore and Coller, 2021). Thus, 3’ cleavage and polyadenylation are critical to the production of functional proteincoding transcripts.

In humans, cleavage and polyadenylation are carried out by a 7-subunit protein complex known as cleavage and polyadenylation specificity factor (CPSF) (Kumar et al., 2019; Zhang et al., 2020). CPSF is comprised of two stable subcomplexes: mammalian polyadenylation specificity factor (mPSF) and mammalian cleavage factor (mCF). These are equivalent to the polymerase module and nuclease module, respectively, of the yeast cleavage and polyadenylation factor (CPF) (Casañal et al., 2017). mPSF contains four subunits: CPSF160, WDR33, CPSF30 and hFip1 (Schönemann et al., 2014). Structures of mPSF/polymerase module in apo and RNA-bound states have recently been elucidated (Casañal et al., 2017; Clerici et al., 2017, 2018; Sun et al., 2018). These showed how two zinc fingers of CPSF30, along with a surface on WDR33, recognize the hexameric polyadenylation signal (PAS) sequence, most commonly AAUAAA, thereby recruiting CPSF to cleavage sites on pre-mRNAs. The poly(A) polymerase enzyme (PAP) is not a stable subunit of human CPSF, but it is instead recruited to cleaved transcripts by hFip1 (Chan et al., 2014; Kaufmann et al., 2004). mCF consists of three subunits: CPSF73, CPSF100 and symplekin (Zhang et al., 2020). CPSF73 is a zinc-dependent RNA endonuclease that belongs to the metallo-β-lactamase family. CPSF100 is a pseudonuclease that is structurally homologous to CPSF73 (Mandel et al., 2006). mCF is tethered to mPSF through a conserved interaction between CPSF160 and a peptide within CPSF100, known as the mPSF interaction motif (PIM) (Zhang et al., 2020; Rodriguez-Molina et al., 2021).

To ensure that mature transcripts of a correct length are produced, pre-mRNAs must be cleaved at specific sites. Deregulation of this process can result in transcriptional defects and nonfunctional transcripts, and can lead to human disease (Curinha et al 2014). *In vitro,* purified CPSF/CPF is an inherently inactive endonuclease, which presumably must be activated by accessory factors to enable strict regulation of 3’-cleavage (Hill et al., 2019; Mandel et al., 2006; Zhang et al., 2020). For example, cleavage stimulatory factor (CStF) and cleavage factor IIm (CFIIm) are both multi-subunit protein complexes involved in cleavage (Takagaki et al., 1990; De Vries et al., 2000). CStF has been shown to bind a G/U-rich region downstream of the cleavage site on pre-mRNAs and provides specificity for poly(A) site selection (Takagaki and Manley, 1997). Another accessory factor, cleavage factor Im (CFIm) is not essential for 3’-cleavage, but it recruits CPSF to pre-mRNAs containing an upstream UGUA motif and contributes to alternative polyadenylation in human cells (Zhu et al., 2018).

The cleavage activity of human CPSF has been studied by functional genomics and by *in vitro* experiments in fractionated nuclear extracts prepared from cultured human cells (for recent examples see Eaton et al., 2018; Schäfer et al., 2018). However, the full protein composition of partially purified nuclear extract is not known, making it difficult to infer molecular mechanism. Moreover, generating mutants of endogenous proteins to test hypotheses is cumbersome.

To enable more detailed mechanistic studies of CPSF endonuclease activation, an *in vitro* assay containing a well-defined set of highly pure proteins is required. Recently, this has been achieved for the human histone pre-mRNA 3’-end processing complex, which shares the endonuclease subunit CPSF73, but differs from CPSF in most of its other subunits and its mechanism of RNA recognition (Gutierrez et al., 2021; Sun et al., 2020). The endonuclease activity of the budding yeast CPF complex has also been reconstituted from purified recombinant proteins (Hill et al., 2019). The minimal active subcomplex in yeast, called core CPF, contains orthologs of CPSF subunits as well as an additional protein Mpe1. In the accompanying manuscript, we show that Mpe 1 is an essential activator of the CPF endonuclease (Rodriguez-Molina et al., 2021). However, while many aspects of 3’-end processing are conserved, there appear to be some differences between the yeast and human machineries, including in RNA specificity and recognition (Rodriguez-Molina et al., 2021; Tian and Graber, 2012). The human ortholog of Mpe1, RBBP6, has been implicated in pre-mRNA 3’-end processing in humans, but has not been defined as a CPSF subunit (Di Giammartino et al., 2014; Shi et al., 2009). Whether it plays a direct role in the cleavage reaction remains unclear.

Here, we reconstitute CPSF from purified proteins that is active in both cleavage and polyadenylation. We demonstrate that human RBBP6 is required for the activation of 3’-end cleavage, even though it is not a stable subunit of CPSF. Our results show that the mechanism of endonuclease activation by Mpe1/RBBP6 is likely to be highly conserved.

## Results

### CStF, CFIIm and RBBP6 are required for activation of CPSF endonuclease

To gain insight into how the human CPSF endonuclease is activated, we attempted to reconstitute pre-mRNA cleavage activity from purified recombinant proteins. We used baculovirus-mediated expression in insect cells to produce highly pure protein complexes predicted to be directly involved in canonical pre-mRNA 3’-end processing. This included CPSF (assembled from individually purified mPSF and mCF subcomplexes) as well as the accessory factors CStF and CFIIm (Figure 1A). Unstructured regions were removed from the CPSF subunits WDR33 and hFip1, and the CFIIm subunit Pcf11 to facilitate purification (Schäfer et al., 2018; Sun et al., 2018). We hypothesized that the conserved region of the multi-domain protein RBBP6 (residues 1-335; Supplementary Information) might also be required for endonuclease activation. RBBP6 did not co-purify with CPSF and was therefore expressed and purified separately. As a model pre-mRNA substrate, we used a 218-nt fragment of the SV40 pre-mRNA, which has been shown to be cleaved efficiently *in vivo* (Kwon et al., 2020; Ryner and Manley, 1987). We omitted the PAP enzyme and ATP from the reactions to focus on the cleavage step of pre-mRNA 3’-end processing (Figure 1B).

**Figure 1.**
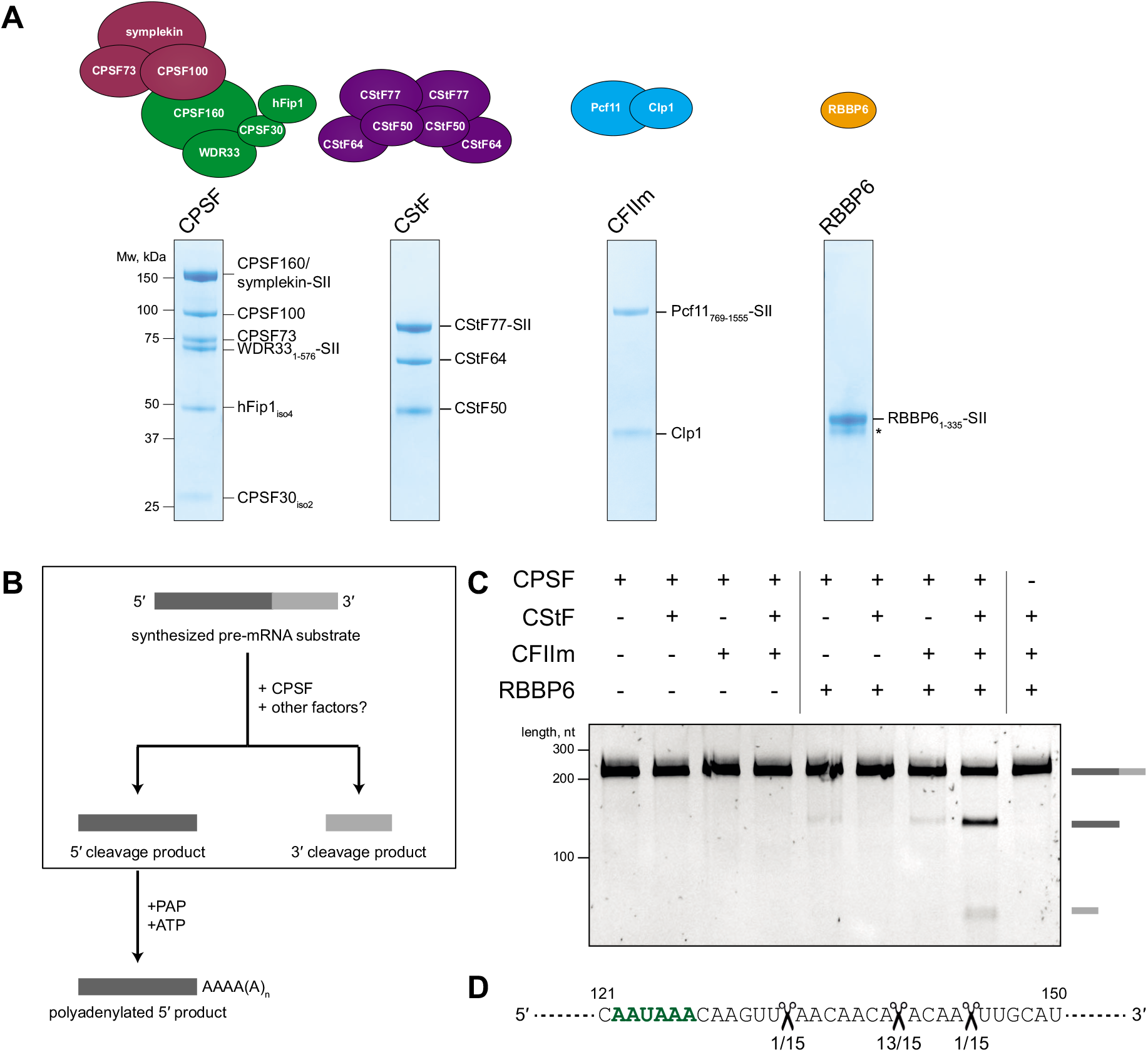
CStF, CFIIm and RBBP6 are required for the activation of CPSF endonuclease. (A) Schematic representations and SDS-PAGE analyses of the purified proteins used in the *in vitro* endonuclease assays. Residue boundaries and alternative isoforms are indicated for truncated proteins. Asterisk denotes degradation products. SII – StrepII tag. (B) Schematic representation of the *in vitro* pre-mRNA 3’-end processing assay. The cleavage reaction is boxed out. The polyadenylation step was not assayed here. (C) Denaturing gel electrophoresis of the SV40 pre-mRNA substrate after incubation with various combinations of human 3’-end processing factors. The full length and cleaved RNAs are shown schematically on the right. (D) Part of the sequence of the SV40 pre-mRNA substrate with the CPSF cleavage sites indicated (scissors). The frequency of a particular cleavage site identified by sequencing of 15 cleavage products is shown below. The polyadenylation signal (PAS) sequence is marked in green. See also Figures S1 and S2.

We tested various combinations of 3’-end processing factors in cleavage assays and analyzed the results by denaturing gel electrophoresis of RNA (Figure 1C). No cleavage activity was observed when the SV40 pre-mRNA was incubated with CPSF alone. Addition of CStF and CFIIm, either individually or together, failed to activate CPSF. However, addition of RBBP6 activated CPSF in the presence of CStF and CFIIm, promoting efficient cleavage of the pre-mRNA substrate. Previous assays in nuclear extract used molecular crowding agents such as polyvinyl alcohol (Adamson et al., 2005), but these were not required here. Omitting CPSF from the reaction did not lead to substrate cleavage, showing that the observed endonuclease activity cannot be attributed to potential contaminants that co-purify with the accessory proteins. Overall, we determined that activation of the CPSF endonuclease requires three additional protein factors: CStF, CFIIm and RBBP6.

Since the SV40 pre-mRNA contains an upstream UGUA motif, we tested whether CFIm affected cleavage of the SV40 substrate with purified CPSF. CFIm is known to bind the RDE domain of hFip1, which is lacking in isoform 4 of hFip1 (Zhu et al., 2018). Therefore, we purified CPSF containing the full-length hFip1 subunit (Figure S1A). Addition of CFIm into the cleavage assay did not provide any further stimulation to CPSF endonuclease in our reconstituted system (Figure S1B). Nevertheless, CFIm may affect cleavage in other conditions or on substrates with multiple UGUA motifs and/or multiple potential PASs.

PAP is dispensable for CPSF cleavage activity, highlighting the fact that cleavage and polyadenylation can be uncoupled *in vitro* (Moore and Sharp, 1985; Ryner and Manley, 1987). Addition of PAP and ATP into a cleavage assay resulted in the polyadenylation of the 5’ cleavage product with heterogeneous poly(A) tail lengths (Figure S1C and D). Thus, our reconstituted CPSF complex is active in both cleavage and polyadenylation.

To identify the precise CPSF cleavage site on the SV40 pre-mRNA substrate, we sequenced several 5’ cleavage products. This revealed that the majority (13/15) of cleaved RNAs were cut 13 nt downstream of the PAS within a CA|A motif, where | indicates the cleavage site (Figure 1D) (Sheets et al., 1987). This is consistent with the known sequence preference of 3’-endonucleases and with previous observations that pre-mRNAs in cells are cleaved 10-30 nt downstream of the PAS (Beaudoing et al., 2000; Hill et al., 2019).

We also tested whether recombinant CPSF could cleave a different pre-mRNA substrate. Under the same reaction conditions, the adenoviral L3 pre-mRNA was cut with similar efficiency as the SV40 pre-mRNA (Figure S2A), suggesting that the same complement of accessory proteins (CStF, CFIIm, RBBP6) are required for activation of the CPSF endonuclease on multiple different pre-mRNA substrates.

### Pre-mRNA cleavage by purified, recombinant CPSF is dependent on CPSF73 and a PAS

Next, we aimed to understand the specificity of the reconstituted 3’-cleavage reaction. First, we generated a CPSF complex containing an active site mutant of CPSF73 (D75N H76A), in which the coordination of catalytic zinc ions was disrupted (Sun et al., 2020). The complex with a mutant endonuclease subunit was inactive in a cleavage assay, suggesting that the observed endonuclease activity is attributable to CPSF73 (Figures 2A and S2B).

**Figure 2.**
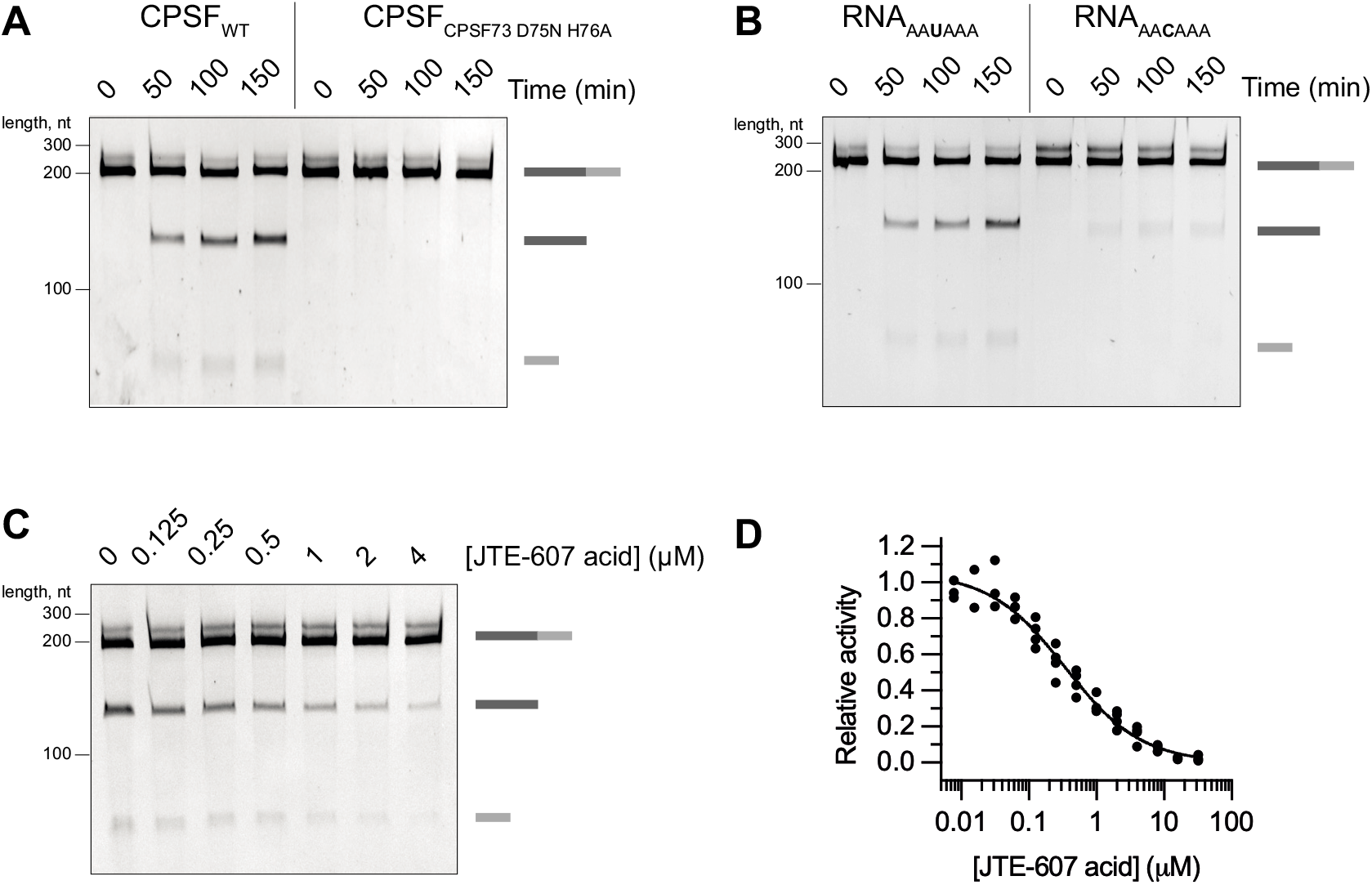
Purified recombinant CPSF pre-mRNA cleavage is dependent on CPSF73 and a PAS. (A) Time-course cleavage assays of the SV40 pre-mRNA substrate comparing the activities of wild-type (CPSF_WT_) and nuclease-dead (CPSF_CPSF73 D75N H76A_) CPSF complexes. (B) Time-course cleavage assays of SV40 pre-mRNA substrates containing either a canonical PAS (RNA_AAUAAA_) or a mutant PAS (RNA_AACAAA_) sequence. (C) Cleavage assays in the presence of increasing concentrations of the JTE-607 acid compound. (D) Dose-response curve of the CPSF cleavage activity as a function of the concentration of JTE-607 acid. Each dot represents a single measurement. See also Figures S2 and S3.

We tested the activity of the mCF subcomplex alone and found that it is completely inactive in the absence of mPSF (Figure S3). The CPSF30 and WDR33 subunits within mPSF recognize the PAS sequence and contribute to specific recruitment of CPSF73 to pre-mRNAs (Clerici et al., 2018; Sun et al., 2018). Replacement of the canonical AAUAAA polyadenylation signal in the SV40 pre-mRNA with an AACAAA hexamer resulted in a loss of cleavage by CPSF, demonstrating that CPSF specifically cleaves PAS-containing RNAs (Figure 2B). It is likely that mPSF is not only involved in RNA binding but is also required for conformational rearrangements that allow endonuclease activation (Rodriguez-Molina et al., 2021).

Recently, CPSF73 was identified as the direct target of JTE-607 – a prodrug with antiinflammatory and anti-cancer properties (Kakegawa et al., 2019; Ross et al., 2020). The active acid form of JTE-607 inhibits the purified, recombinant yeast 3’-endonuclease (Ross et al., 2020) and also inhibits CPSF73 within the human histone pre-mRNA 3’-end processing complex *in vitro* (Gutierrez et al., 2021). We therefore tested whether the JTE-607 acid analog was also inhibitory to the reconstituted human canonical 3’-end processing complex. Titrating the compound into the cleavage reaction showed dose-dependent inhibition of the endonuclease activity with an IC50 of ~350 nM (Figure 2C-D), which is very similar to the *Kd* of the acid form of JTE-607 for isolated CPSF73 (~370 nM; Ross et al 2020). This confirms that the observed *in vitro* endonuclease activity is specific to CPSF73.

### Canonical and histone pre-mRNA 3’-endprocessing complexes are activated by different mechanisms

The human histone pre-mRNA 3’-processing reaction was recently reconstituted with purified proteins, and the structure of the substrate-bound complex was determined in an active state (Sun et al 2020; Gutierrez et al 2021). The histone processing complex shares three subunits with CPSF: symplekin, CPSF100 and CPSF73 (termed the histone cleavage complex, equivalent to mCF in CPSF). Although some aspects of endonuclease activation are carried out by proteins exclusive to the histone complex, the N-terminal domain (NTD) of symplekin (which is also found in CPSF) was shown to be essential for activating CPSF73. We tested whether the symplekin NTD plays a similar role in CPSF. To this end, we prepared a CPSF complex in which the NTD of symplekin was deleted. The CPSF complex lacking the symplekin NTD retained a similar activity as wild-type CPSF, suggesting that the mechanism of endonuclease activation is not conserved between the two CPSF73-containing complexes (Figure 3A).

**Figure 3.**
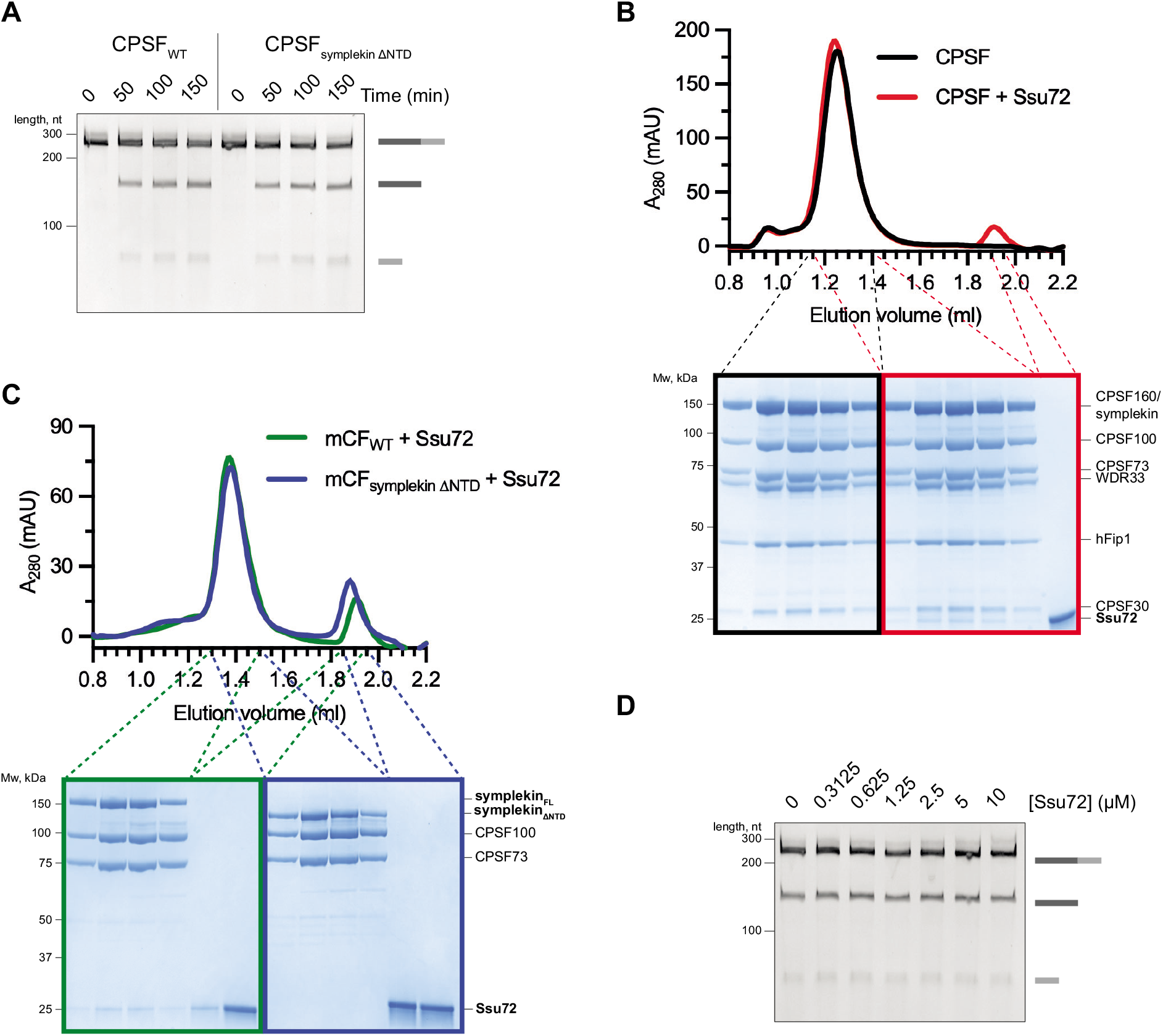
Canonical and histone pre-mRNA 3’-end processing complexes are activated by different mechanisms. (A) Time-course cleavage assays of the SV40 pre-mRNA substrate comparing wild-type CPSF (CPSF_WT_) and CPSF lacking the symplekin NTD (CPSF_symplekin δNTD_). (B) Gel filtration chromatograms (top) and SDS PAGE analyses (bottom) of CPSF in the presence or absence of Ssu72. (C) Gel filtration chromatograms (top) and SDS PAGE analyses (bottom) of wild-type mCF (mCF_WT_) and mCF lacking the NTD of symplekin (mCF_symplekin δNTD_), mixed with Ssu72. (D) Cleavage assays in the presence of increasing concentrations of Ssu72.

In addition, the phosphatase Ssu72 was shown to inhibit the histone processing complex by binding to and sequestering the symplekin NTD (Sun et al 2020). Ssu72 is a subunit of yeast CPF (Casañal et al., 2017), and hence we tested whether it also interacts with human CPSF. We found that Ssu72 interacts with mCF and CPSF but not with mCF lacking the symplekin NTD (Figure 3B-C). However, titrating Ssu72 into the CPSF cleavage reaction did not affect the *in vitro* endonuclease activity (Figure 3D). Together, these results suggest that, similar to the histone 3’-processing complex, Ssu72 interacts with the symplekin NTD in CPSF but the mechanism of CPSF73 activation is fundamentally different in each complex.

### RBBP6 is a conserved activator of canonical pre-mRNA 3 ‘-end cleavage

We were particularly intrigued by the role of RBBP6 in activating the CPSF endonuclease. Human RBBP6 is an ~200 kDa protein with a conserved N-terminal region containing several ordered domains, and a long, disordered, non-conserved C-terminal tail, which interacts with various binding partners not directly related to mRNA 3’-end processing (Batra et al., 2018; Li et al., 2007, Sakai et al., 1995; Simons et al., 1997) (Figure 4A and Supplementary Information). A construct encompassing only the N-terminal domains of RBBP6 was sufficient to stimulate CPSF (Figure 1C), suggesting that the C-terminal region is dispensable for pre-mRNA cleavage *in vitro*.

**Figure 4.**
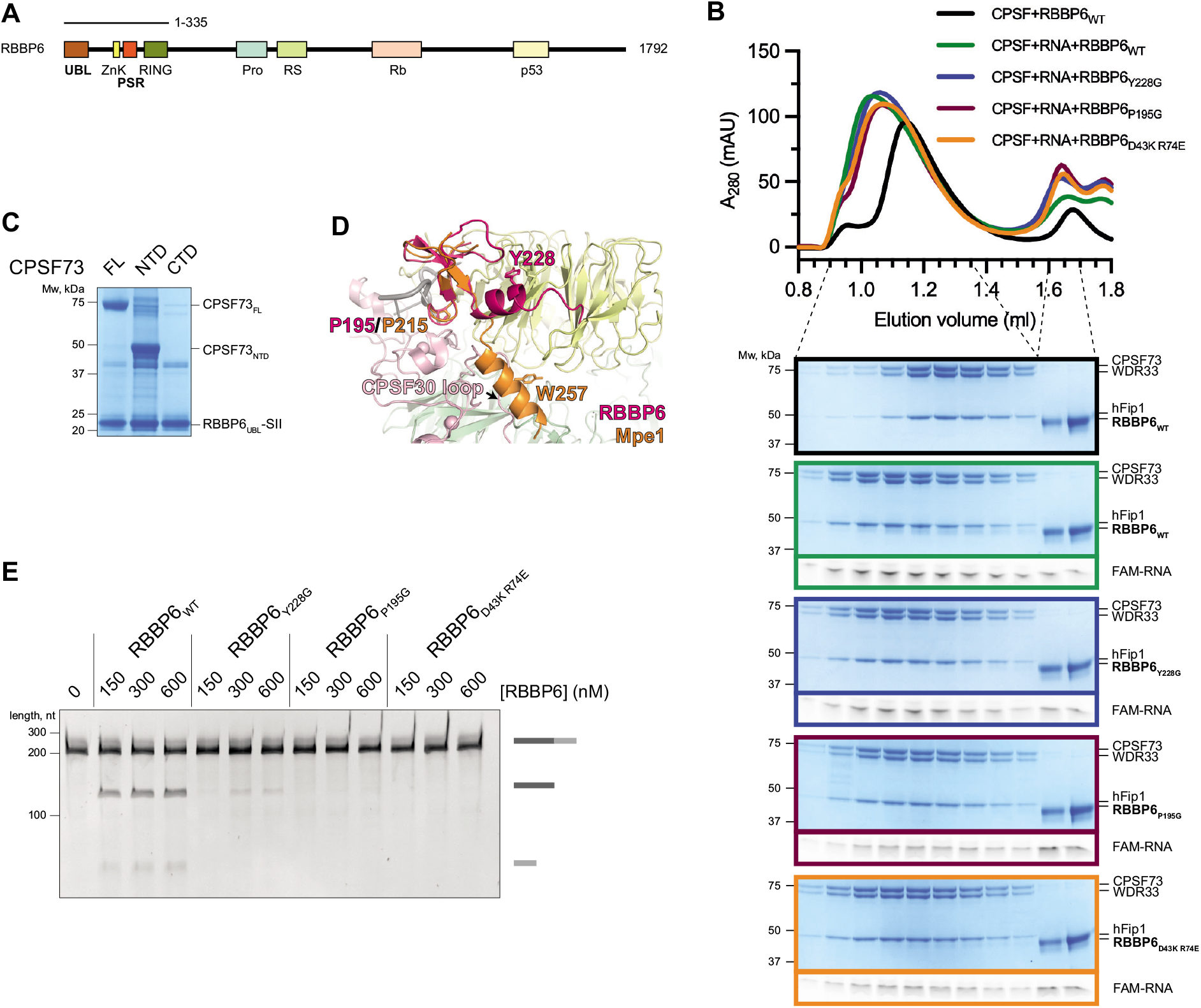
RBBP6 is a conserved activator of canonical pre-mRNA 3’-end cleavage. (A) Domain diagram of full-length human RBBP6 (1,792 residues). The construct used in this study (residues 1-335) is indicated. UBL – ubiquitin-like domain; ZnK – zinc knuckle; PSR – PAS-sensing region; Pro – proline-rich domain; RS – arginine, serine-rich domain; Rb – retinoblastoma protein-interacting region; p53 – p53-interacting region. (B) Gel filtration chromatograms of CPSF and RBBP6 in the presence or absence of a 5’-FAM fluorescently-labelled 41 nt L3 RNA (top), and denaturing PAGE analysis of proteins and RNA from the indicated fractions (bottom). The gels are cropped and outlined in color to correspond with the colors of the chromatogram traces. (C) Pull-down of SII-tagged UBL domain of RBBP6 in the presence of various constructs of CPSF73 from Sf9 insect cells. RBBP6 pulls-down FL and NTD CPSF73, but not CTD CPSF73. FL – full-length; NTD – N-terminal domain (residues 1-460); CTD – C-terminal domain (residues 461-684). (D) Overlay of the experimental structure of the yeast Mpe1 PSR (orange; Rodriguez-Molina et al 2021) and an AlphaFold2 prediction of the structure of the equivalent region in human RBBP6 (magenta) overlaid on human mPSF (Sun et al, 2018). Residues of functional significance are indicated. A loop of CPSF30 would clash with the C-terminal helix of the Mpe1 PSR. Yellow – WDR33; pink – CPSF30; green – CPSF160; grey – PAS RNA. (E) Cleavage assays in the presence of various concentrations of either wild-type (RBBP6_WT_) or mutant (RBBP6_Y228G_/RBBP6_P195G_/RBBP6_D43K R74E_) RBBP6. See also Figures S4 and S5.

The yeast ortholog of RBBP6, Mpe1, is a constitutive subunit of yeast CPF (Casañal et al., 2017; Vo et al., 2000). In contrast, RBBP6 does not co-purify with either endogenous or recombinant CPSF complexes (Chan et al., 2014). To investigate whether RBBP6 binds stably to CPSF, we used size exclusion chromatography. When mixed together, RBBP6 and CPSF eluted from the column in two separate peaks, indicating that any potential interaction between them does not survive over the column (Figure 4B). However, when a 41-nt fragment of L3 pre-mRNA was included, a substoichiometric amount of RBBP6 co-migrated with RNA-bound CPSF (Figure 4B). We also performed pull-downs using MS2-tagged L3 pre-mRNA and found that RBBP6 was pulled down by RNA only in the presence of CPSF (Figure S4A). This suggests that RBBP6 is recruited transiently to CPSF in an RNA-dependent manner, which is reminiscent of RNA-mediated stabilization of Mpe1 on the yeast polymerase module (Rodriguez-Molina et al 2021).

Yeast Mpe1 contacts two subunits of CPF (Hill et al 2019; Rodriguez-Molina et al 2021). First, the ubiquitin-like domain (UBL) of Mpe1 stably interacts with the N-terminal nuclease domain (NTD) of the endonuclease subunit (Hill et al 2019). The interacting residues are highly conserved and a structure of the complex could be confidently modelled (Hill et al 2019). An isoform of RBBP6 that contains only the UBL domain inhibits cleavage in nuclear extract by competing with full-length RBBP6 (Di Giammartino et al., 2014).

To test whether human RBBP6 and CPSF73 interact in a similar manner to the yeast proteins, we co-expressed StrepII-tagged RBBP6-UBL with various constructs of CPSF73 in insect cells and performed pull-down studies. This showed that tagged RBBP6-UBL pulled down stoichiometric amounts of both full-length CPSF73 and the CPSF73 nuclease domain (NTD), but it did not interact with the CPSF73 C-terminal domain (CTD) (Figure 4C). Thus, the interaction of RBBP6-UBL and the CPSF73 nuclease is conserved in humans. However, the complex between RBBP6-UBL and the CPSF73 nuclease domain dissociated during anion exchange chromatography, demonstrating that the affinity between the human proteins is relatively low, explaining why a stable complex was not observed on gel filtration (Figure 4B). We introduced mutations in RBBP6-UBL (D43K and R74E) at the putative RBBP6-CPSF73 interaction interface (Hill et al., 2019). The RBBP6-D43K-R74E mutant failed to activate CPSF in a cleavage assay and no longer associated with the CPSF-RNA complex (Figure 4B and 4E). These results highlight that, like in yeast, the RBBP6-UBL contacts the CPSF73 nuclease domain.

In the accompanying manuscript, we show that yeast Mpe1 binds to the Pfs2 subunit of the yeast polymerase module and directly contacts the pre-mRNA substrate (Rodriguez-Molina et al 2021). We therefore named this region of Mpe1 the PAS-sensing region (PSR). The PSR sequence is conserved in RBBP6, and using AlphaFold2 (Jumper et al., 2021), we predict that this region is likely to adopt a similar overall structure to the Mpe1 PSR (Figures 4D and Supplementary Information). In the predicted structure, the C-terminal helix of the RBBP6 PSR is in an alternative binding position on WDR33. Interestingly, the site of Mpe1 interaction is occupied by a loop of CPSF30 in the human complex.

To test the functional relevance of the RBBP6 PSR, we mutated a conserved aromatic residue in RBBP6, Y228. This residue is equivalent to W257 in Mpe1, which forms critical contacts with the yeast polymerase module. We also mutated P195, which contacts RNA in the yeast complex (Rodriguez-Molina et al 2021). Both RBBP6-Y228G and RBBP6-P195G mutants were almost completely ineffective at activating the CPSF endonuclease (Figure 4E). In addition, neither RBBP6 mutant co-migrated with CPSF during gel filtration chromatography, even in the presence of RNA (Figure 4B). Since RBBP6 and Mpe1 have been implicated in RNA binding on their own (Baejen et al., 2014; Lee and Moore, 2014), we compared the relative affinities of the RBBP6 mutants for RNA using electrophoretic mobility shift assays (EMSAs). None of the mutations affected the ability of RBBP6 to bind the RNA used in gel filtration assays (Figure S4B), which suggests that the mutated residues are involved in binding CPSF directly, not in binding RNA. These observations demonstrate that the PSR of RBBP6 plays a crucial role in stimulating the endonuclease.

Together, these data suggest that RBBP6 interacts transiently with CPSF in an RNA-dependent manner to act as an essential activator of the canonical pre-mRNA 3’-endonuclease. RBBP6 interactions with CPSF and, therefore the mechanism of endonuclease activation by RBBP6, are likely to be conserved from yeast to human.

## Discussion

3’-cleavage of nascent protein-coding transcripts is essential for both mRNA maturation and transcription termination. Here, we reconstitute the canonical pre-mRNA 3’-endonuclease activity of human CPSF with purified proteins and determine that CStF, CFIIm and RBBP6 are all required for activation. Together, these four factors likely represent the minimal and universal machinery that cleaves pre-mRNAs at their 3’-ends. In agreement with this, orthologous factors (core CPF and CF IA) are required in yeast (Hill et al, 2019). However, yeast CF IB is also required to enforce specificity of cleavage. There is no clear ortholog of CF IB in humans.

Purified CPSF73 in isolation only weakly and non-specifically cleaves RNA (Mandel et al., 2006). Thus, its incorporation into a 7-subunit protein complex may ensure that the endonuclease is inhibited until it is specifically activated on PAS-containing transcripts. The additional requirement for three RNA-binding accessory factors would further restrict activation, precisely positioning the endonuclease on RNA and preventing premature cleavage. *In vivo*, variations in nuclear concentrations of basal cleavage factors (as has been shown for CStF) (Takagaki and Manley, 1998) as well as other accessory proteins (for example, CFIm) (Tseng et al., 2021) additionally regulate cleavage site selection in a transcript- and context-specific manner (Gruber and Zavolan, 2019). It has been proposed that human CPSF100 may also be able to catalyze endonucleolytic cleavage (Kolev et al., 2008). However, under the conditions used here, CPSF73 is the only active endonuclease within CPSF.

Previously, RBBP6 was suggested to regulate mRNA 3’-end site selection (Di Giammartino et al., 2014), but its role has been largely underestimated, primarily because RBBP6 is not a constitutive subunit of human CPSF. In contrast, yeast Mpe1 is a core subunit of CPF (Casañal et al., 2017; Hill et al., 2019; Vo et al., 2000). Despite differences in affinity, the molecular nature of the interaction of RBBP6/Mpe1 with CPSF/CPF is likely conserved, as demonstrated in our mutational analyses. The affinities of other components of the 3’-end processing machinery also differ between human and yeast. For example, the poly(A) polymerase enzyme is a constitutive subunit of the yeast but not the human complex (Chan et al., 2014; Kaufmann et al., 2004). Human CPSF has a nanomolar affinity for PAS-containing RNA (Hamilton et al., 2019), while the interaction of CPF with RNA is orders of magnitude weaker (Hill et al 2019). In addition, human CStF and CFIIm are separate complexes, whereas in yeast, they form a constitutive complex called CF IA (Gordon et al., 2011; Schäfer et al., 2018). These differences may enable alternative types of regulation of pre-mRNA 3’-end processing in different organisms, whilst retaining the same fundamental mechanism of endonucleolytic cleavage.

RBBP6 interacts with CPSF in an RNA-dependent manner. This RNA dependence explains why RBBP6 was detected in an RNA-bound post-cleavage 3’-end processing complex but not in endogenous apo CPSF (Chan et al., 2014; Shi et al., 2009). The C-terminal domain of RBBP6 is absent from our construct, and it is not required for cleavage *in vitro*. Interestingly, this domain contains linear peptide motifs that bind transcription factors (Rb and p53) (Saijo et al., 1995; Sakai et al., 1995; Simons et al., 1997), and it also has an RS domain, which in other proteins is known to bind the spliceosome and SR proteins that regulate alternative splicing (Graveley and Maniatis, 1998). Therefore, RBBP6 may coordinate 3’-end processing with transcription and splicing *in vivo*.

The *in vitro* endonuclease activity of human CPSF is substantially slower than that of yeast CPF under similar conditions (Hill et al., 2019; Rodriguez-Molina et al., 2021). This could be due to the transient nature of RBBP6 binding to CPSF or because additional, unknown protein factors are involved *in vivo.* However, it is also possible that human CPSF is an inherently inefficient and potentially more accurate endonuclease to allow more extensive regulation, for example, to enable correct cleavage site selection even on very long 3’-UTRs with multiple potential PASs (Martin et al., 2012). On the other hand, CPF cleavage must be very efficient to prevent transcriptional readthrough into downstream genes in yeast, where genes are closely spaced (David et al., 2006; Rodriguez-Molina et al., 2021).

The structure of the active histone pre-mRNA 3’-end processing machinery demonstrated how the propagation of conformational rearrangements across many protein factors can lead to the opening of the CPSF73 active site (Sun et al., 2020). Although we show that the precise nature of CPSF73 activation differs between CPSF and histone complexes, we envision that a coordinated assembly of CPSF, CStF, CFIIm and RBBP6 on a pre-mRNA substrate leads to a similar conformational change in CPSF73.

CPSF73-specific inhibitors have been demonstrated to have anti-cancer (Kakegawa et al 2019; Ross et al 2020), and anti-protozoan (Jacobs et al. 2011; Palencia et al. 2017; Sonoiki et al. 2017; Swale et al. 2019) properties. Thus, although the mechanisms of mRNA 3’-end processing are highly conserved, the differences between species can be exploited. The reconstitution of human canonical pre-mRNA 3’-end processing with purified proteins provides new opportunities for studying the molecular mechanism of CPSF in detail both biochemically and structurally.

## Data Availability

All gel files will be uploaded to a data repository.

## Author Contributions

V.B. designed and performed experiments, analyzed data and wrote the manuscript. T.E. synthesized JTE607. J.C, supervised the research. L.A.P. supervised the research, analyzed data and wrote the manuscript.

## Acknowledgements

We thank Ana Casañal for assistance with construct design, Manuel Carminati for help with protein purification, James Stowell for help with RNA sequencing, Max Wilkinson for MBP-MS2 protein, and Juan Rodriguez-Molina, Ananth Kumar and other members of the Passmore lab for helpful discussions and advice. This work was supported by the Medical Research Council, as part of United Kingdom Research and Innovation, MRC file reference numbers MC_U105192715 (L.A.P.), MC_U105181009 (J.C.) and MC_UP_A024_1008 (J.C.); the European Union’s Horizon 2020 research and innovation programme (ERC Consolidator grant agreement 725685) (to L.A.P) and a Herchel Smith PhD Studentship from the University of Cambridge (to V.B.).

## MATERIALS AND METHODS

### Cloning

#### CPSF, CStF, CFIIm, CFIm, RBBP6, PAP, Ssu72

*E. coli* codon-optimised genes encoding each full-length protein and isoform 2 of CPSF30 (Uniprot O95639-2) in pACEBac vectors were synthesized by Epoch Life Science. All cloning was validated by sequencing (Source Bioscience).

To generate isoform 4 of hFip1 (Uniprot Q6UN15-4), fragments containing residues 1-28 and 44-393 were amplified by PCR. Substitution F393K was also introduced during the PCR of fragment 44-393. Both fragments were assembled into an empty pACEBac vector using Gibson assembly.

To express Ssu72 in *E. coli,* the coding region of Ssu72 was amplified by PCR from its pACEBac vector. The forward primer contained an NdeI cleavage site, and the reverse primer had a BamHI cleavage site. After digestion with NdeI (NEB, cat. No. R0111) and BamHI-HF (NEB, cat. No. R3136) enzymes, the Ssu72 coding region was ligated into an empty pET-28a vector that had been cleaved with the same enzymes. The vector contained an in-frame His6-tag followed by a 3C protease cleavage site on its 5’-end.

The full coding regions of CStF77, symplekin, CFIm25 and PAP were amplified by PCR from their original pACEBac vectors and cloned using Gibson assembly into pACEBac vectors containing an in-frame TEV cleavage site followed by an SII-tag on its 3’-end. For the following genes, only the sequences encoding the indicated residues were amplified by PCR: 1-576 of WDR33, 769-1555 of Pcf11, 1-335 of RBBP6, 1-142 of RBBP6 (RBBP6_UBL_), 341-1274 of symplekin (symplekin_△NTD_). These fragments were also cloned into pACEBac-TEV-SII vectors as described above.

To produce catalytically inactive CPSF73 D75N H76A, the CPSF73 pACEBac plasmid was divided into three overlapping fragments and these fragments were amplified by PCR. The mutations were located in the overlapping region between two of the three fragments. All three fragments were ligated together using Gibson assembly. To produce CPSF73_NTD_ and CPSF73_CTD_ constructs, CPSF73 residues 1-460 and 461-684, respectively, were amplified by PCR and assembled into empty pACEBac vectors using Gibson assembly.

#### Assembly into pBigl vectors

A modified biGBac protocol was used to generate pBig 1 vectors encoding all subunits of each complex (mPSF, mCF, CStF, CFIIm, CFIm and their variants) as described previously (Hill et al., 2019; Weissmann et al., 2016).

### Protein expression

#### Baculovirus

pBig1 (mPSF, mCF, CStF, CFIIm, CFIm) or pACEBac (RBBP6, PAP) vectors were transformed into EMBacY cells. Extracted bacmids were transfected into Sf9 insect cells to generate P1 virus. To produce P2 virus, Sf9 cells were infected with P1 virus. Proteins were overexpressed by infecting large-scale cultures of Sf9 cells (except for mPSF, which was overexpressed in Hi5 insect cells) with P2 virus. The cells were harvested by centrifugation when the cell viability fell below ~90% (after 3-4 days). The cell pellets were flash-frozen in liquid N_2_ and stored at −80°C. All these procedures were described in detail previously (Hill et al., 2019; Kumar et al., 2021).

#### E. coli

*E. coli* BL21(DE3) Star cells transformed with the His6-Ssu72-encoding vector were induced with 0.5 mM IPTG at OD_600_ ~0.6 and grown overnight at 20°C. The cells were harvested by centrifugation, flash-frozen in liquid N_2_ and stored at −80°C.

### Protein purification

#### mPSF-hFipl_iso4_

A frozen cell pellet of Hi5 cells was thawed in lysis buffer (50 mM HEPES-NaOH pH 8.0, 300 mM NaCl, 1 mM TCEP, 2 mM Mg(OAc)_2_), supplemented with 50 μg/ml DNaseI, 3 protease inhibitor tablets (Roche, cat. No. 11836153001) and 1 ml BioLock (IBA, cat. No. 20205-050) per 1 l cell culture. The cells were lysed by sonication, and the lysate was cleared by centrifugation. The cleared lysate was filtered through a 0.65 μm filter and incubated with Strep-Tactin beads (IBA, cat. No. 2-1201-025) for 2-3 h. The beads were washed with lysis buffer, and the complex was eluted with 2.5 mg/ml desthiobiotin (IBA, cat. No. 2-1000-005) in lysis buffer. The eluate was diluted to reduce the NaCl concentration to 75 mM, filtered through a 0.45 μm filter and applied to a 1 ml Resource Q column (Cytiva, cat. No. 17117701) equilibrated in buffer A (20 mM HEPES-NaOH pH 8.0, 75 mM NaCl, 0.5 mM TCEP, 2 mM Mg(OAc)_2_). The complex was eluted using a linear gradient of buffer B (20 mM HEPES-NaOH pH 8.0, 1 M NaCl, 0.5 mM TCEP, 2mM Mg(OAc)_2_) over 50 column volumes. The peak fractions were pooled, concentrated and injected onto a Superose 6 XK 17/600 pg column (Cytiva, cat No. 71501695) equilibrated in size exclusion buffer (20 mM HEPES-NaOH pH 8.0, 150 M NaCl, 0.5 mM TCEP, 2 mM Mg(OAc)_2_). Selected fractions were pooled and concentrated. The concentrated protein was aliquoted, flash-frozen in liquid N_2_ and stored at −80°C.

#### mPSF-hFip1_FL_

mPSF-hFip1_FL_ was purified from Hi5 cells by Strep-Tactin affinity chromatography and anion exchange chromatography as described for mPSF-hFip1_iso4_. The peak fractions of mPSF-hFip1_FL_ from a 1 ml Resource Q column were pooled, aliquoted, flash-frozen in liquid N_2_ and stored at −80°C.

#### mCF, mCF_CPSF73 D75NH76A_, mCF_symplekin ΔNTD_

mCF and its variants were purified from Sf9 cells using the same protocol as mPSF but: 1) 50 μg/ml RNaseA was added to lysis buffer; 2) buffers were supplemented with 5% v/v glycerol before each concentration step; 3) size exclusion buffer contained 20 mM HEPES-NaOH pH 8.0, 150 M NaCl, 1 mM TCEP.

#### CStF

CStF was purified from Sf9 cells using the same protocol as mCF, except that the size exclusion buffer contained 20 mM HEPES-NaOH pH 8.0, 200 mM NaCl, 1 mM TCEP.

#### CFIIm

CFIIm was purified from Sf9 cells using the same protocol as mCF with a few modifications. In the lysis buffer, DNaseI and RNaseA were replaced by 50 U/ml benzonase (Merck, cat. No. E1014), and 100 μM PMSF (Merck, cat. No. 93482) was also added. The size exclusion buffer of CFIIm contained 20 mM Tris-HCl pH 8.5, 150 M NaCl, 0.5 mM TCEP, 5% v/v glycerol.

#### RBBP6, RBBP6_Y228G_, RBBP6_PI95G_, RBBP6_D43KR74E_

RBBP6 was purified from Sf9 cells using the same protocol as mPSF but with different buffers: Lysis buffer – 50 mM HEPES-NaOH pH 8.0, 400 mM NaCl, 1 mM TCEP; buffer A – 20 mM HEPES-NaOH pH 8.0, 40 mM NaCl, 0.5 mM TCEP; buffer B – 20 mM HEPES-NaOH pH 8.0, 1 M NaCl, 0.5 mM TCEP; size exclusion buffer – 20 mM HEPES-NaOH pH 8.0, 200 mM NaCl, 0.5 mM TCEP, 2 mM Mg(OAc)_2_. Also, HiLoad 16/600 Superdex 200 pg column (Cytiva, cat. No. 28989335) was used for the size exclusion step.

#### CFIm

CFIm was purified from Sf9 cells by Strep-Tactin affinity chromatography and anion exchange chromatography as described for mPSF-hFip1_iso4_ but using different buffers: lysis buffer – 50 mM bicine-NaOH pH 9.0, 400 mM NaCl, 0.5 mM TCEP, 2 mM Mg(OAc)_2_, 10% v/v glycerol; buffer A – 20 mM bicine-NaOH pH 9.0, 150 mM NaCl, 0.5 mM TCEP, 2 mM Mg(OAc)_2_, 10% v/v glycerol; buffer B – 20 mM bicine-NaOH pH 9.0, 1 M NaCl, 0.5 mM TCEP, 2 mM Mg(OAc)_2_, 10% v/v glycerol. The peak fractions of CFIm from a 1 ml Resource Q column were pooled, aliquoted, flash-frozen in liquid N_2_ and stored at −80°C. Before running assays, ~ 100 μl CFIm was thawed and dialyzed overnight against 500 ml dialysis buffer (20 mM bicine-NaOH pH 9.0, 400 mM NaCl, 0.5 mM TCEP, 2 mM Mg(OAc)_2_, 10% v/v glycerol).

#### PAP

PAP was purified from Sf9 cells by Strep-Tactin affinity chromatography as described for mPSF-hFip1_iso4_. The eluate was incubated overnight at 4°C with 20 μg/ml TEV protease to remove the StrepII tag. The protein was further purified using a 1 ml HiTrap Q column (Cytiva, cat. No. 29051325) equilibrated in buffer A (50 mM HEPES-NaOH pH 8.0, 100 mM NaCl, 1 mM TCEP) and eluted with a linear gradient of buffer B (50 mM HEPES-NaOH pH 8.0, 1 M NaCl, 1 mM TCEP). The peak fractions were concentrated and loaded onto a HiLoad 26/600 Superdex 200 pg column (Cytiva, cat. No. 28989336) equilibrated in buffer containing 50 mM HEPES-NaOH pH 8.0, 150 mM NaCl, 1 mM TCEP. The peak fractions were pooled, concentrated, and aliquoted. The aliquots were flash-frozen in liquid N_2_ and stored at −80°C.

#### Ssu72

Cells were lysed by sonication in buffer A (50 mM HEPES-NaOH pH 8.0, 500 mM NaCl, 1 mM TCEP, 20 mM imidazole) supplemented with 2 protease inhibitor tablets and 50 μg/ml DNaseI. The lysate was cleared by centrifugation and loaded onto a HisTrap HP 5 ml column (Cytiva, cat. No. 17524701) equilibrated in buffer A. The protein was eluted with a linear gradient of buffer B (50 mM HEPES-NaOH pH 8.0, 500 mM NaCl, 1 mM TCEP, 500 mM imidazole) over 20 column volumes. 43 μg/ml 3C protease was added to the pooled peak fractions to remove the His6-tag, and the protein was dialyzed overnight using a 7 kDa-cut-off membrane against dialysis buffer (50 mM HEPES-NaOH pH 8.0, 500 mM NaCl, 1 mM DTT). The dialyzed sample was concentrated with 5% v/v glycerol and loaded onto a HiLoad Superdex 75 26/600 column (Cytiva, cat. No. 28989334) equilibrated in size exclusion buffer (20 mM HEPES-NaOH pH 8.0, 200 mM NaCl, 1 mM TCEP). The peak fractions were concentrated in the presence of 5% v/v glycerol, aliquoted and flash-frozen in liquid nitrogen. The protein was stored at −80°C.

### Preparation of RNA substrates

5’-FAM fluorescently-labelled 41 nt L3 RNA was synthesized by IDT.

The DNA sequences encoding fragments of SV40 pre-mRNA with either wild-type (AAUAAA) or mutant PAS (AACAAA) were purchased as gBlocks from IDT. The sequence of the T7 RNA polymerase promoter was added to the 5’-end of the gBlock by PCR amplification.

The template of the L3 pre-mRNA was purchased from IDT as a gBlock. The fragment had a KpnI (NEB, cat. No. R0142) cleavage site on its 5’-end and a BamHI site on its 3’-end. After restriction digest with both enzymes, the L3 fragment was ligated into a linearized pUCIDT plasmid encoding the T7 RNA polymerase promoter followed by three MS2 loops upstream of the insert.

#### In vitro transcription

All pre-mRNA substrates were transcribed using HiScribe T7 High Yield RNA Synthesis Kit (NEB, cat. No. E2040) and subsequently purified with Monarch RNA Cleanup Kit (NEB, cat. No. T2040).

### Cleavage assays with purified proteins

Each protein factor was first diluted in protein dilution buffer (20 mM HEPES-NaOH pH 7.25 (measured at room temperature), 150 mM NaCl, 0.5 mM TCEP, 2 Mg(OAc)_2_). Individually purified mPSF and mCF complexes were mixed at 2.5 μM each in protein dilution buffer and incubated on ice for 30 min. All protein components were then mixed on ice in 19 μl per condition and/or time point at the final concentrations of 50 nM CPSF, 100 nM CStF, 100 nM CFIIm and 300 nM RBBP6 in a buffer containing 20 mM HEPES-NaOH pH 7.25 (measured at room temperature), 50 mM NaCl, 0.5 mM TCEP, 2 Mg(OAc)_2_ and 1 U/μl RiboLock (Thermo, cat. No. EO0381). The tubes were transferred to 37°C, and the reaction was initiated by an addition of the RNA substrate to a final concentration of 100 nM. Unless indicated otherwise, the reactions were stopped after 150 min by adding 5 μl stop buffer (130 mM EDTA, 5% v/v SDS, 12 mg/ml proteinase K in protein dilution buffer) and incubating them at 37°C for a further 15 min. The samples were mixed with RNA Gel Loading Dye (Thermo Scientific, cat. No. R0641) and loaded on a pre-run (30 W for 30 min) denaturing 10% (SV40) or 6% (L3) polyacrylamide gel containing 7 M urea in TBE buffer. The gels were run for 25 min at 400 V, stained with SYBR Green (Invitrogen, cat. No. S7564) and imaged using a ChemiDoc XRS+ (BioRad).

### Coupled cleavage and polyadenylation assay with purified proteins

3 μM mPSF-hFip1_iso4_, 3 μM mCF and 6 μM PAP were mixed and loaded onto a Superose 6 Increase 3.2/300 column (Cytiva, cat No. 29091598) equilibrated in HEPES-NaOH pH 8.0, 150 mM NaCl, 0.5 mM TCEP, 2 mM Mg(OAc)_2_. The peak fraction was added to a cleavage reaction either without or with 2 mM ATP. The assays were run and the results analyzed as described above.

### Sequencing of 5’-cleavage products

A standard cleavage reaction of the SV40 substrate was analyzed on a denaturing gel as described above. The band corresponding to the 5’-cleavage product was excised and submerged in 50 μl crush and soak buffer (3 M Na(OAc) pH 5.2, 0.1 M EDTA pH 7.4, 20% v/v SDS). The gel band was crushed with a sterile pipette tip and incubated overnight at 37°C. After taking off the supernatant, the same steps were repeated with 50 μl fresh crush and soak buffer for 2 h. The two supernatants were combined, and the extracted RNA was precipitated at −20°C for 2 h in 300 μl absolute ethanol with 1μl Glycoblue (Invitrogen, cat. No. AM9516). The RNA was pelleted in a chilled microcentrifuge at maximum speed for 10 min and washed with 500 μl 70% ethanol. The RNA pellet was resuspended in 20 μl DEPC water. An adenylated adaptor of known sequence was ligated to the 3’-end of the extracted 5’-cleavage product using T4 RNA ligase 2, truncated (NEB, cat. No. M0242). The RNA was purified from the ligation reaction components using Monarch RNA Cleanup Kit. The 5’-cleavage products that contained the adaptor were converted into cDNA using SuperScript IV First-Strand Synthesis System (Invitrogen, cat. No. 18091050) with a forward primer specific to a 5’-region of the SV40 RNA and a reverse primer that anneals to the adaptor. The cDNA was further amplified by PCR and ligated into a bacterial vector using Zero Blunt PCR Cloning Kit (Invitrogen, cat. No. K270040). After transformation into TOP10 *E. coli* cells, 15 colonies were picked, and the isolated plasmids were sequenced using the M13R primer (Source Bioscience) to determine the 3’-end of the 5’-cleavage product.

### Assays with JTE-607 acid compound

#### Synthesis

The prodrug of JTE-607 was purchased from Tocris and hydrolyzed to JTE-607 acid analog as previously described (Ross et al., 2020).

#### Titration of JTE-607 acid compound

Standard cleavage assays were set up in the presence of various concentrations of the acid form of JTE-607, and the samples were analyzed by denaturing polyacrylamide gel electrophoresis as described above. The relative activity of CPSF at each concentration of the drug was calculated as the relative intensity of the cleavage product bands in each lane relative to this ratio in the absence of the drug (x – JTE-607 concentration):

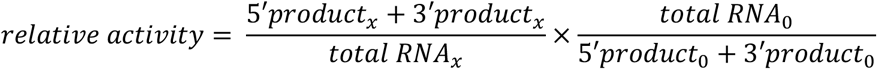

The intensity values were measured in Fiji. The data were plotted in Prism 9 and fitted to the equation of “[Inhibitor] vs. response – Variable slope (four parameters)” with an R^2^ value of 0.9656.

### Pull-downs from insect cells

A P2 virus encoding RBBP6_UBL_-SII and a P2 virus carrying a gene of one of the CPSF73 variants (CPSF73_FL_, CPSF73_NTD_, CPSF73_CTD_) were used to co-infect Sf9 cells at ~2 million cells/ml. The cultures were harvested after 3 days by centrifugation and washed in ice-cold PBS. The cell pellets were lysed using glass beads (Merck, cat. No. G8772) in lysis buffer (50 mM HEPES-NaOH pH 8.0, 300 mM NaCl, 1 mM TCEP, 2 mM Mg(OAc)_2_) supplemented with 2 protease inhibitor tablets per 50 ml buffer. The lysates were cleared by centrifugation and applied to Strep-Tactin beads. After a 2 h incubation, the beads were washed in lysis buffer, and the bound proteins were eluted by incubating the samples in NuPAGE LDS Sample Buffer (Invitrogen, cat. No. NP0007) at 98°C for 2 min. The eluted proteins were analyzed on a NuPAGE 4-12% Bis-Tris 1.0 mm Mini Protein Gel (Invitrogen, cat. No. NP0321) and stained with Instant Blue (Abcam, cat. No. 119211).

### Gel filtration chromatography

All samples were incubated on ice for 30 min before analysis. To investigate RBBP6 binding to CPSF, 2.5 μM CPSF and 7.5 μM RBBP6 or its point mutants were mixed with or without 5 μM 5’-FAM 41-nt L3 RNA. The CPSF-RBBP6 samples were loaded onto a Superose 6 Increase 3.2/300 column (Cytiva, cat No. 29091598) equilibrated in HEPES-NaOH pH 8.0, 50 mM NaCl, 0.5 mM TCEP. To test Ssu72 binding to CPSF and mCF variants, 2.5 μM CPSF/mCF/mCF_sympiekin δNTD_ was incubated with 10 μM Ssu72. The samples were loaded onto the same column but in buffer containing HEPES-NaOH pH 8.0, 150 mM NaCl, 0.5 mM TCEP. The protein content of the peak fractions was analyzed by SDS-PAGE as described above. To detect the RNA, stop buffer was added to an aliquot of each fraction. After incubation at 37°C for 10 min, RNA loading dye was added, and the samples were loaded onto 15% Novex TBE-Urea gels (300 V, 50 min). The gels were scanned using a FAM channel on a Typhoon FLA 7000 instrument (GE Healthcare).

### *In vitro* pull-downs on M2-L3 pre-mRNA

The pull-downs were performed in pull-down buffer containing 20 mM HEPES-NaOH pH 8.0, 50 mM NaCl, 0.5 mM TCEP, 2 mM Mg(OAc)_2_. First, 520-nt MS-L3 pre-mRNA was incubated with MBP-tagged MS2 protein at a molar ratio 1:3 for 45 min on ice. 3 μM RBBP6, 1 μM CPSF or 3 μM RBBP6 + 1μM CPSF were then added and incubated for 1.5 h. The mixture containing RBBP6/CPSF/RBBP6+CPSF and MBP-MS2-bound L3 pre-mRNA was mixed with amylose beads (NEB, cat. No. E8021) equilibrated in pull-down buffer and incubated rotating at 4°C for 1.5 h. The beads were washed with pull-down buffer. Protein-RNA complexes were eluted in pull-down buffer supplemented with 20 mM maltose (Merck, cat. No. 63418). The eluates were loaded on a NuPAGE 4-12% Bis-Tris 1.0 mm Mini Protein Gel. The proteins were transferred onto a nitrocellulose membrane using Trans-Blot Turbo Transfer System (Bio-Rad, cat. No. 1704158). StrepII-tagged proteins (RBBP6, symplekin and WDR33) were detected using streptavidin-HRP conjugate (Merck Millipore, cat. No. 18152) and Amershan ECL Detection Reagents (Cytiva, cat. No. RPN2106). The blots were visualized using a ChemiDoc XRS+ (BioRad).

### Electrophoretic mobility shift assays (EMSAs)

Indicated concentrations of various point mutants of RBBP6 were mixed with 100 nM 41-nt 5’-FAM-labelled L3 pre-mRNA and orange G loading dye (0.4% w/v orange G, 50% v/v glycerol, 1 mM EDTA). The protein-RNA mixtures were incubated on ice for 15 min and then loaded onto a 10% native polyacrylamide gel. The gel was run for 50 min at 100 V at 4°C. The RNA was visualized using a FAM channel on a Typhoon FLA 7000 instrument (GE Healthcare).

---

**Figure S1.**
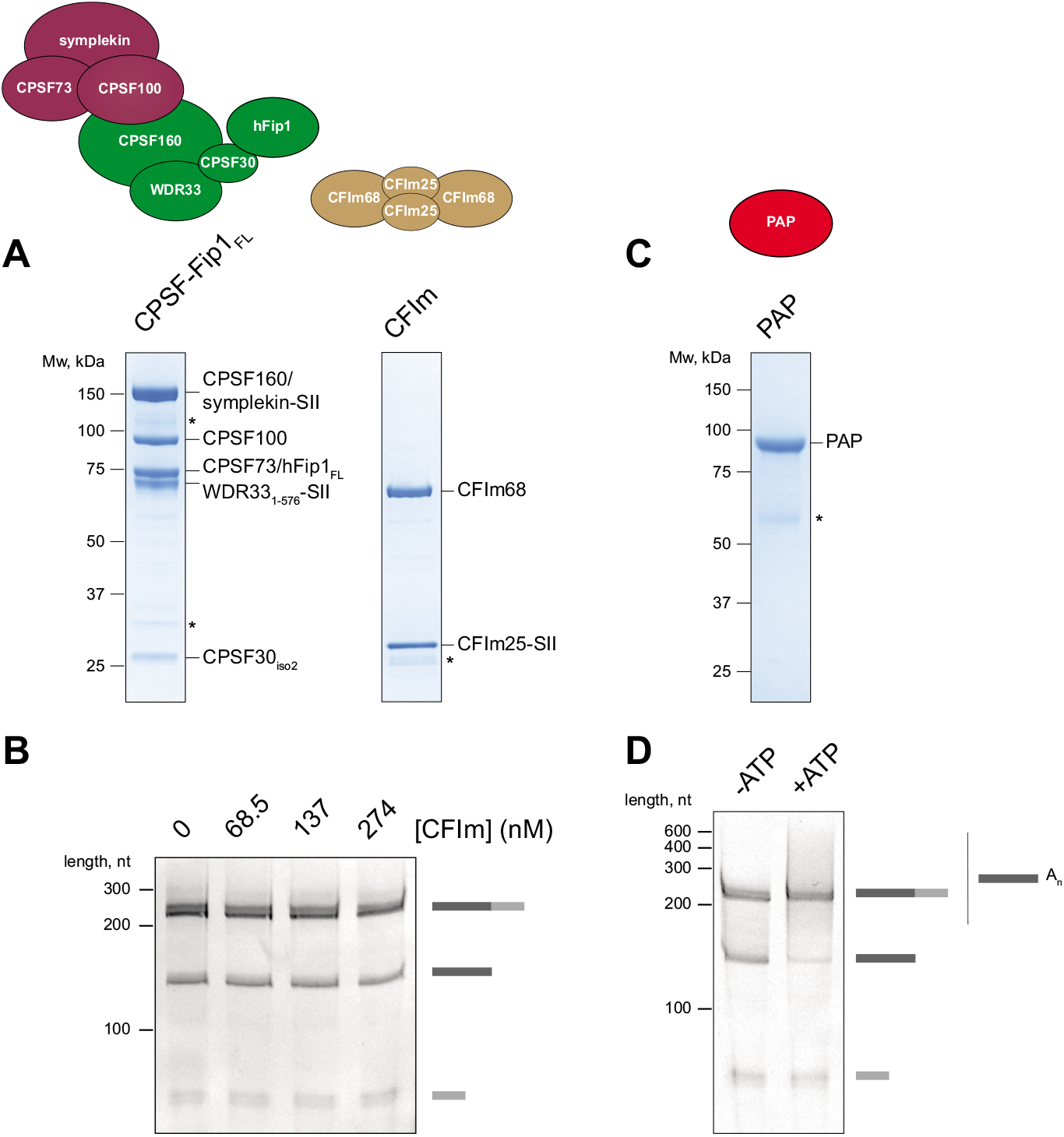
CFIm and PAP are not required for CPSF cleavage activity. Related to Figure 1. (A) SDS-PAGE analyses of purified CPSF containing full-length hFip1 (hFip1_FL_) subunit and of purified CFIm complex. Asterisks denote degradation products. SII – StrepII tag. (B) Cleavage assays of the SV40 pre-mRNA substrate with CPSF-hFip1_FL_ in the presence of increasing concentrations of CFIm. CFIm does not substantially affect CPSF cleavage activity. (C) SDS-PAGE analysis of purified PAP. Asterisk denotes degradation products. (D) Coupled cleavage and polyadenylation assay of the SV40 pre-mRNA substrate in the presence of PAP either without (cleavage only) or with (cleavage and polyadenylation) ATP. Upon addition of ATP, the 5’ cleavage product band disappears and is replaced by a smear corresponding to polyadenylated 5’ products containing poly(A) tails of variable length.

**Figure S2.**
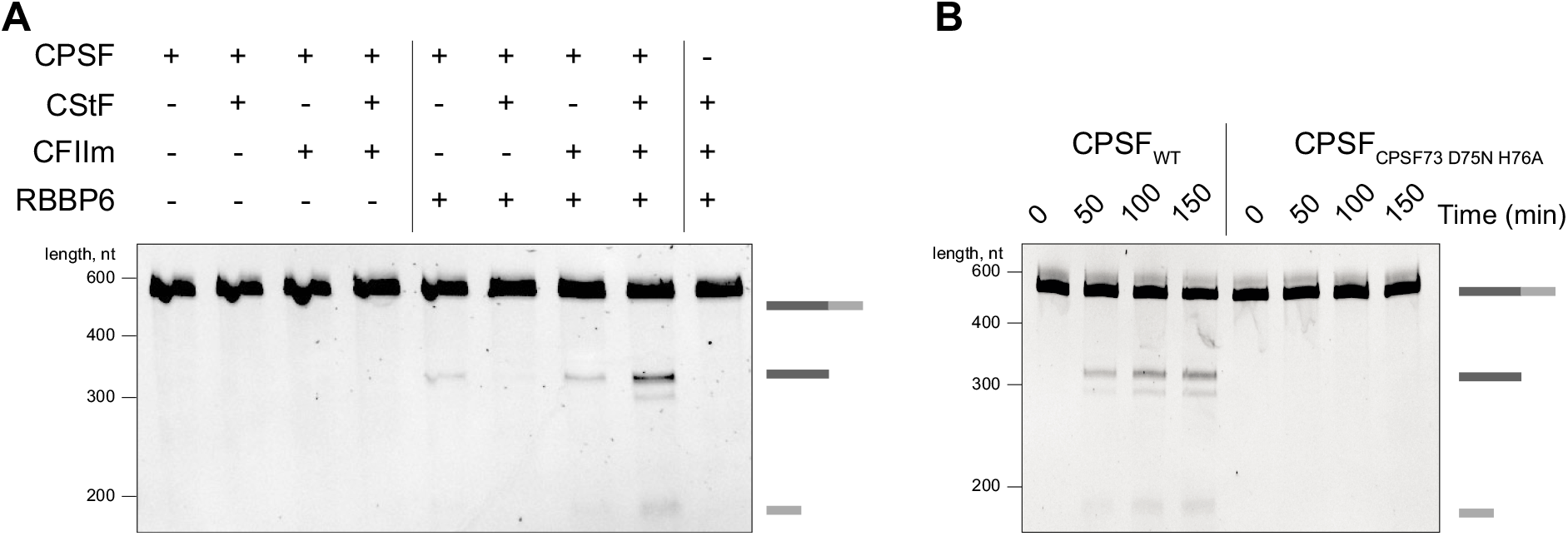
L3 pre-mRNA substrate is cleaved by purified CPSF. Related to Figures 1 and 2. (A) Denaturing gel electrophoresis of the MS2-tagged L3 520-nt pre-mRNA substrate after incubation with various combinations of human 3’-end processing factors. For both L3 and SV40 (Figure 1C), there is a small amount of cleavage by CPSF + RBBP6 and CPSF + RBBP6 + CFIIm but the CPSF complex is substantially activated with RBBP6, CFIIm and CStF. (B) Time-course cleavage assays of the MS2-L3 pre-mRNA substrate comparing wild-type (CPSF_WT_) and nuclease-dead (CPSF_CPSF73 D75N H76A_) CPSF complexes.

**Figure S3.**
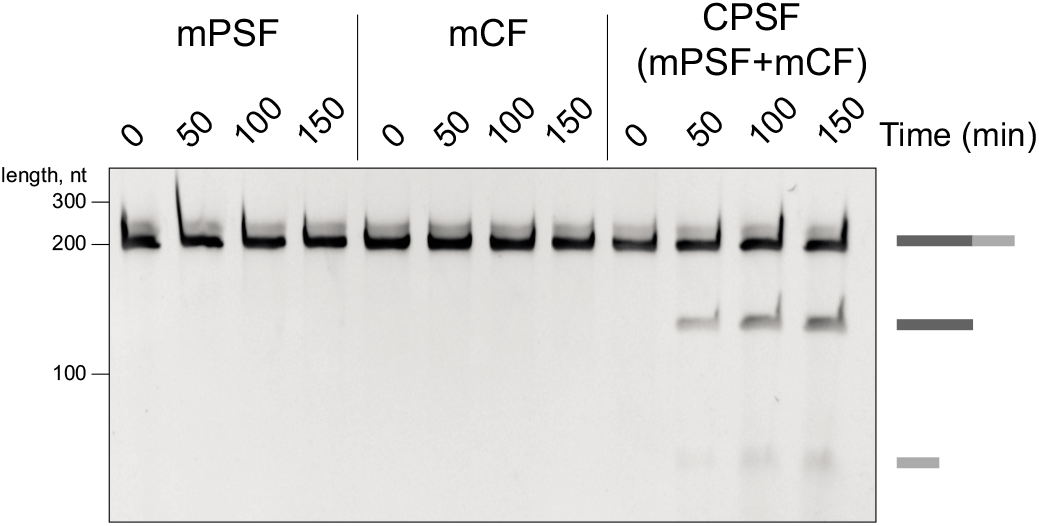
CPSF73 is inactive in the absence of mPSF. Related to Figure 2. Time-course cleavage assays of the SV40 pre-mRNA substrate comparing activities of mPSF, mCF and CPSF (mPSF and mCF combined).

**Figure S4.**
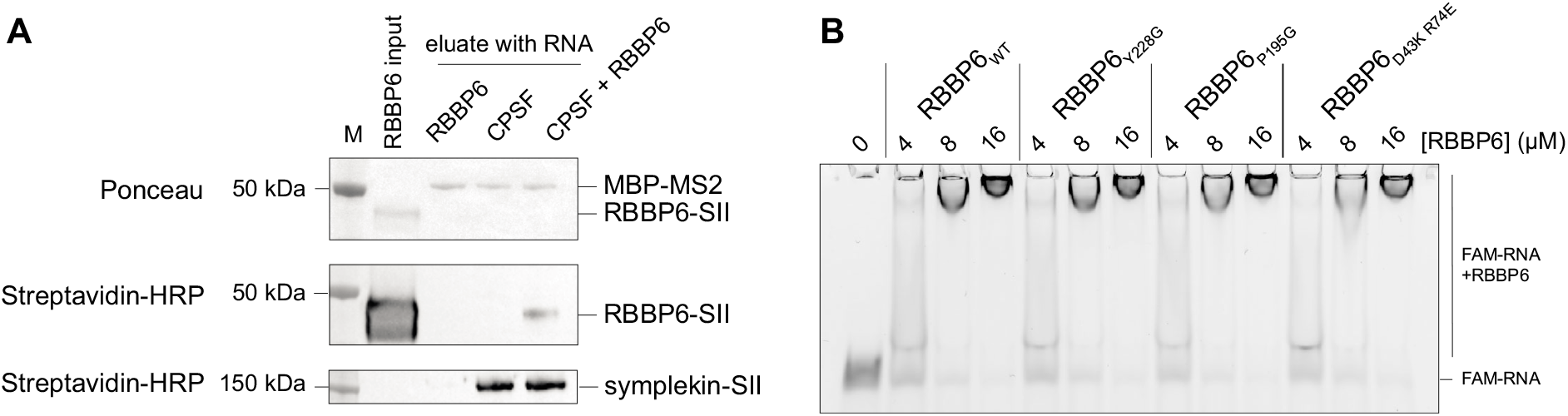
CPSF recruits RBBP6 to pre-mRNAs. Related to Figure 4. (A) *In vitro* pull-downs of RBBP6 by MS2-tagged 520-nt L3 pre-mRNA substrate in the presence and absence of CPSF. The RNA was mixed with MBP-tagged MS2 and the indicated proteins and immobilized on amylose beads. The eluates were analyzed by Western blots against the StrepII tag. Under these experimental conditions, RBBP6 is recruited to RNA only in the presence of CPSF. M – protein molecular weight marker. (B) Electrophoretic mobility shift assays (EMSAs) of the binding to a 5’-FAM fluorescently-labelled 41 nt L3 RNA by RBBP6 and its point mutants. The mutations do not affect RBBP6 binding to RNA in the absence of CPSF.

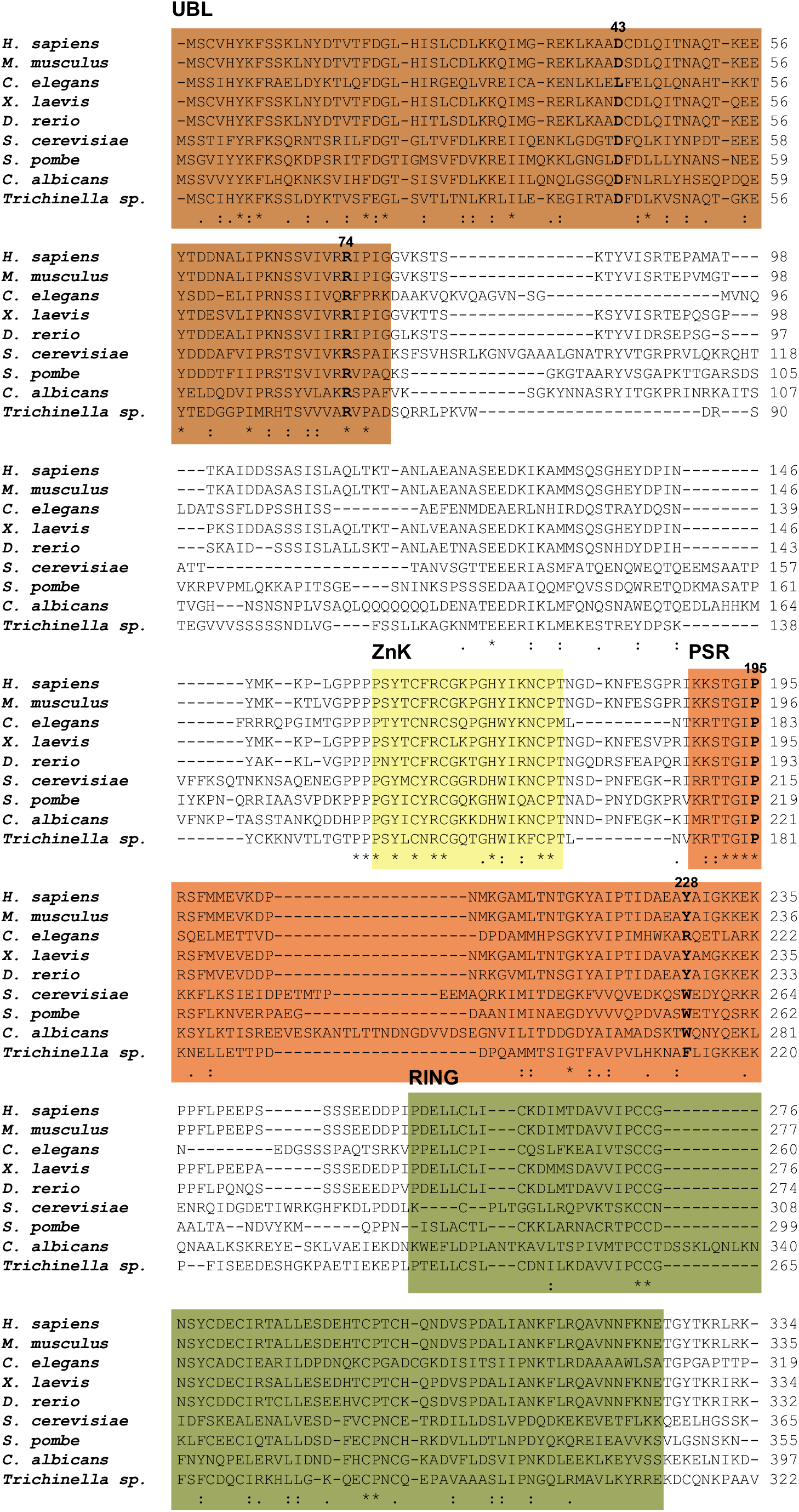

**Supplementary Information. Sequence alignment of RBBP6 orthologs**

Residues corresponding to human RBBP6 1-335 were aligned to orthologs from other eukaryotes. Residues investigated in this study are indicated. UBL – ubiquitin-like domain; ZnK – zinc knuckle; PSR – PAS-sensing region. * – conserved amino acid identity;: – amino acid properties conserved;. – properties partly conserved. The alignment was generated with Uniprot.

